# 3D Printed Nerve Guidance Conduit for Biologics-Free Nerve Regeneration and Vascular Integration

**DOI:** 10.1101/2025.04.30.651603

**Authors:** Jacob Schimelman, David B. Berry, Susie Johnson, Ruskin Shi, Sophie Brown, Quyen T. Nguyen, Shaochen Chen

## Abstract

There is a clinical need for an effective nerve guidance conduit to treat peripheral nerve injuries. Many studies have explored different materials and active cues to guide neural regeneration, with some success. However, none have demonstrated a comparable or better functional recovery than the clinical standard autograft. Autografts are often insufficient for reconstruction of an injury to long nerves such as the sciatic or brachial plexus. Synthetic nerve guidance conduits (NGCs) have been investigated for these injuries to guide axonal regeneration and lead to functional recovery. We have designed a biologics-free hydrogel-based multi-channel conduit with defined microscale features to guide axonal outgrowth. To investigate extraneural vascular infiltration and its effects on functional recovery, we also designed a multi-microchannel conduit with defined regularly spaced micropores, orthogonal to the axon guidance channels. Using our custom-built Rapid Projection, Image-guided, Dynamic (RaPID) bioprinting system, we are able to fabricate each hydrogel conduit within minutes from a milliliter-volume prepolymer vat. With our state-of-the-art printing platform, we have achieved NGCs with a consistent channel wall width of 10 microns. We implanted the NGCs for 17 weeks in a murine sciatic nerve transection injury model. We assessed the functional recovery by dynamic gait analysis throughout the recovery period and by compound muscle action potential (CMAP) electrophysiology before NGC harvesting. Both the non-porous and micro-porous conduit groups led to functional nerve regeneration on par with the autograft group. Further, both conduit groups resulted in restoration of bulk motor function to pre-injury performance.

**Translational Impact Statement:** We have developed a synthetic therapeutic that can be implanted to aid nerve regeneration after a debilitating injury that performs similarly to the current clinical standard treatment. One major hurdle to clinically translating an implantable therapeutic is needing to include biologics such as cells or active biomolecules, which increases the regulatory burden. With this in mind, we systematically developed a biologics-free synthetic hydrogel nerve guidance conduit, composed of polymer backbones that have been used in many FDA-approved applications, capable of restoring motor function.

## INTRODUCTION

Peripheral nerve injuries (PNIs) can occur in everyday life, e.g., from motor vehicle accidents; workplace accidents related to heavy machinery or power tools usage; and in combat-related blast injuries, e.g., from IEDs. In the US alone, there are over 800,000 PNI-related surgical operations performed annually ^1^. If left unrepaired, a PNI can result in chronic pain and/or paralysis. The current standard of care for reconstructive management of PNI the clinical setting is a sensory nerve autograft ^2,3^. Though autografts have been useful during reconstructive surgery, they are limited by donor site morbidity ^4^. Additionally, they are constrained by a potential mismatch between the length of the defect site and donor nerve. For major injuries to the brachial plexus or the sciatic nerves, the two longest sets of peripheral nerves in the body, using autografts may not be a viable approach.

Due to the above surgical considerations, there is intense interest in developing a synthetic nerve guidance conduit to replace the need for an autograft ^5^. For over 20 years, there have been numerous studies published on investigating cylindrical polymeric conduits, with or without inner topological guidance, either by themselves or combined with a growth factor drug delivery treatment and/or cellular therapy ^5–8^. Yet, none have resulted in an FDA-approved system that has replaced the autograft as the standard of care for the surgical management of PNI. What has been approved are synthetic polymer hollow tubes or wraps to bridge the gap distance of the defect site, but they do not provide sufficient topological guidance or bioactive cues ^1,9,10^. There has been progress on commercializing decellularized human nerve allografts as an off-the-shelf alternative ^1^, but this will not fully meet the need of replacing autografts due to the inherent supply limitations of human cadaver donated nerves.

We have approached developing a compelling replacement to the autograft by systematically designing a nerve conduit to provide the minimum features necessary to lead to equivalent or better performance for motor function recovery in comparison to treatment by the reverse autograft. It has been well-established that microstructure provides topological guidance to the extending axon ^11–27^. Additionally, it has been shown that the stiffness and mechanical integrity of a conduit has a significant effect on the regeneration quality ^1,28^.

Our previous NGC had four circular-cross-section microchannels that encouraged tissue regeneration across the length of the conduit of nerve bundles with multiple fascicles of axons ^28^. The conduits were produced via our custom Rapid Projection, Image-guided, Dynamic (RaPID) 3D-printing system (Figure 1 A). We demonstrated that a photopolymerized composite hydrogel of poly(ethylene glycol) diacrylate (PEGDA) and gelatin methacryloyl (GelMA) was biocompatible within the murine sciatic nerve injury environment, with minimal to no evidence of an adverse inflammatory response. Additionally, there was preliminary evidence suggesting sensorimotor functional recovery by 6 weeks post injury (wpi). From this initial study, a 400-μm-diameter microchannel was shown to successfully guide axonal extension in vivo and the axons physically interacted only partially with the channel’s inner surface area topography.

**Figure 1.**
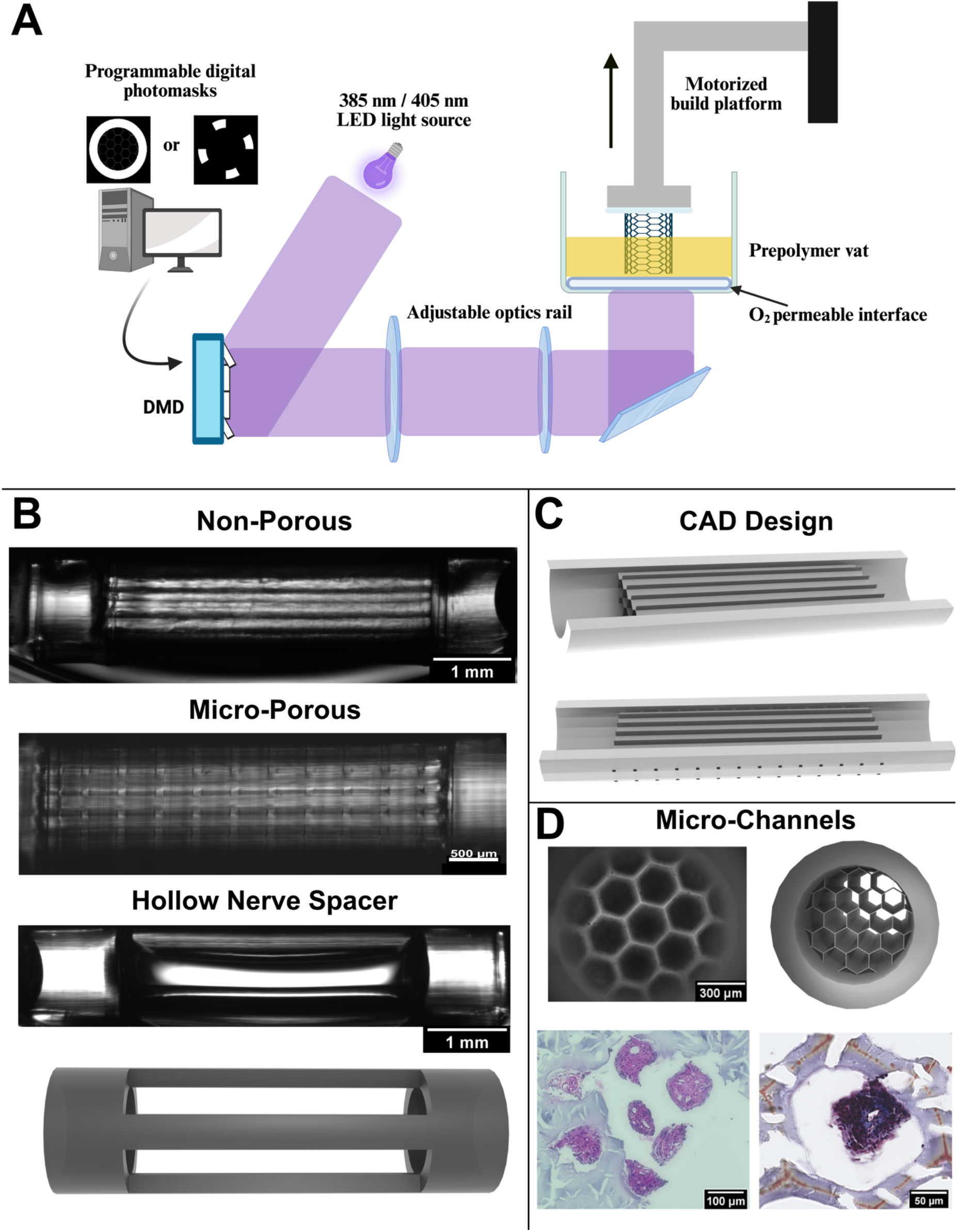
Development of a 3D-printed microchannel nerve guidance conduit. (A) Schematic of the RaPID bioprinting system.^38^ (B) Longitudinal brightfield image of the non-porous nerve guidance conduit, micro-porous, and hollow nerve spacer. Below the hollow nerve spacer image is its corresponding CAD image. (C) Longitudinal cross-sections of the CAD images corresponding to the non-porous (top) and micro-porous (bottom) microchannel conduits. (D) Transverse cross-section brightfield (top left) and corresponding CAD (top right) images are shown. An H&E stained paraffin section (bottom left) showing representative fasciculated axon bundles extending through the seven central microchannels, and a Masson’s Trichrome stained cryosection (right) showing a representative single fascicle extending through one of the central microchannels.

Most reports in the NGC literature have focused on axon guidance alone, but there has been recent interest in developing NGCs capable of enhancing intraneural blood vessel outgrowth within fascicles and/or extraneural blood vessel outgrowth from the distal tissue towards the injury site ^29–33^. The extraneural vasculature is known to be activated by the secreted VEGF-A from the circulating macrophages that are responding to the injury site, resulting in angiogenesis towards the regenerating axons^34^. The blood vessels provide a topological bridge across the injury site for Schwann cell migration, thus injury-induced angiogenesis plays a critical role in innate peripheral nerve regeneration ^34^. However, although blood vessel outgrowth has been shown to be critical for nerve regeneration, it remains unclear how increasing the amount of blood vessels within a NGC affects tissue regeneration, and more critically, functional recovery.

One reason for the lack of translation of NGC research to the clinic is that they are generally fabricated using methods that are either difficult to scale or difficult to eliminate batch-to-batch variation. Additive manufacturing, commonly referred to as 3D printing, provides a highly promising approach to both producing consistently at scale for off-the-shelf NGCs as well as for patient-specific pre-surgical planning designs and point-of-care manufacturing. The FDA has recently acknowledged the unique opportunity of implementing 3D printing solutions in the clinic and is currently working to standardize their assessments^35,36^. Another reason for the lack of clinical translation is that most NGCs reported include biologics such as neurotrophic or growth factors, guidance cues, and/or cellular therapies. Including bioactive materials, especially cells, within a biomaterial device greatly increases the regulatory hurdles (and cost) for commercialization.

In this report, we systematically study developing a NGC capable of axon guidance, vascularization, functional nerve regeneration, and motor function recovery without the inclusion of any active drug or biologic to enable a path for direct clinical translation. Using our RaPID bioprinting system, we 3D print a PEGDA-GelMA composite hydrogel conduit with an array of 300-μm-wide microchannels separated by 10-μm-thin walls, where one conduit design version is functionally non-porous and a second conduit design version has a patterned array of 50-μm-wide micropores that we hypothesize would enable extraneural vascular integration.

## RESULTS

### Nerve Guidance Conduit Design

The conduit is designed for a complete transection sciatic nerve injury in a mouse model. This injury model results in the nerve ends retracting and forming a 4-mm gap defect, as previously established ^28^. As such, the microchannel region of the conduit is designed to be 4-mm long, and each end of the conduit featured a 1-mm long hollow sleeve for securing each nerve end (Figure 1 B, C). The microchannel diameter was optimized to be 300 μm to guide individual fascicles (Figure 1 D). Additionally, to maximize the total number of microchannels for individual fascicle guidance, a tessellated hexagon pattern is chosen as the cross-section geometry to increase packing density as compared to circular geometries and the wall thickness between microchannels was minimized to only 10 μm (Figure 1 D). To introduce micropores, we use CAD software to add a regularly patterned array of micropores, spaced 250 μm apart in each direction, throughout the central conduit microchannel region (Figure 1 B, C bottom). A square cross-section geometry with a side length of 50 μm is used to ensure a sufficient entryway for nascent blood vessels ^37^. The RaPID 3D printing system has readily produced this design change with precision (Figure 1 B).

### Development of a Hollow Nerve Spacer Negative Control

To best evaluate the performance of the non-porous and micro-porous conduits, in addition to the positive control of the clinical standard autograft, a negative control is needed. A conventional option is to surgically perform the complete transection sciatic nerve injury without providing a therapeutic treatment. However, this approach suffers from the inability to control the positioning of the nerve ends during recovery. A simple hollow conduit would provide a direct control for topological guidance of the microchannels, but not a true negative control for the injury model. We have solved this problem by designing and 3D-printing a defined-separation construct consisting of the same hollow sleeves from the conduit design that are joined together by four thin pillars, securing the nerve ends at the same 4-mm gap distance as the therapeutic conduit groups (Figure 1 B, bottom).

### Mechanical Testing

To ensure the Young’s modulus of the conduit matches that of reported murine sciatic nerve tissue^39,40^, which is on the order of magnitude of 1 MPa, we perform both compressive testing to obtain the bulk modulus of the 3D-printed PEGDA-GelMA hydrogel material and nanoindentation testing to obtain the local modulus of each conduit design, with and without micropores, as printed (Figure 2). We find that the bulk modulus is 2.63 ± 0.712 MPa. The local moduli for the non-porous and micro-porous conduits are 1.79 ± 0.95 MPa and 1.09 ± 0.33 MPa, respectively.

**Figure 2.**
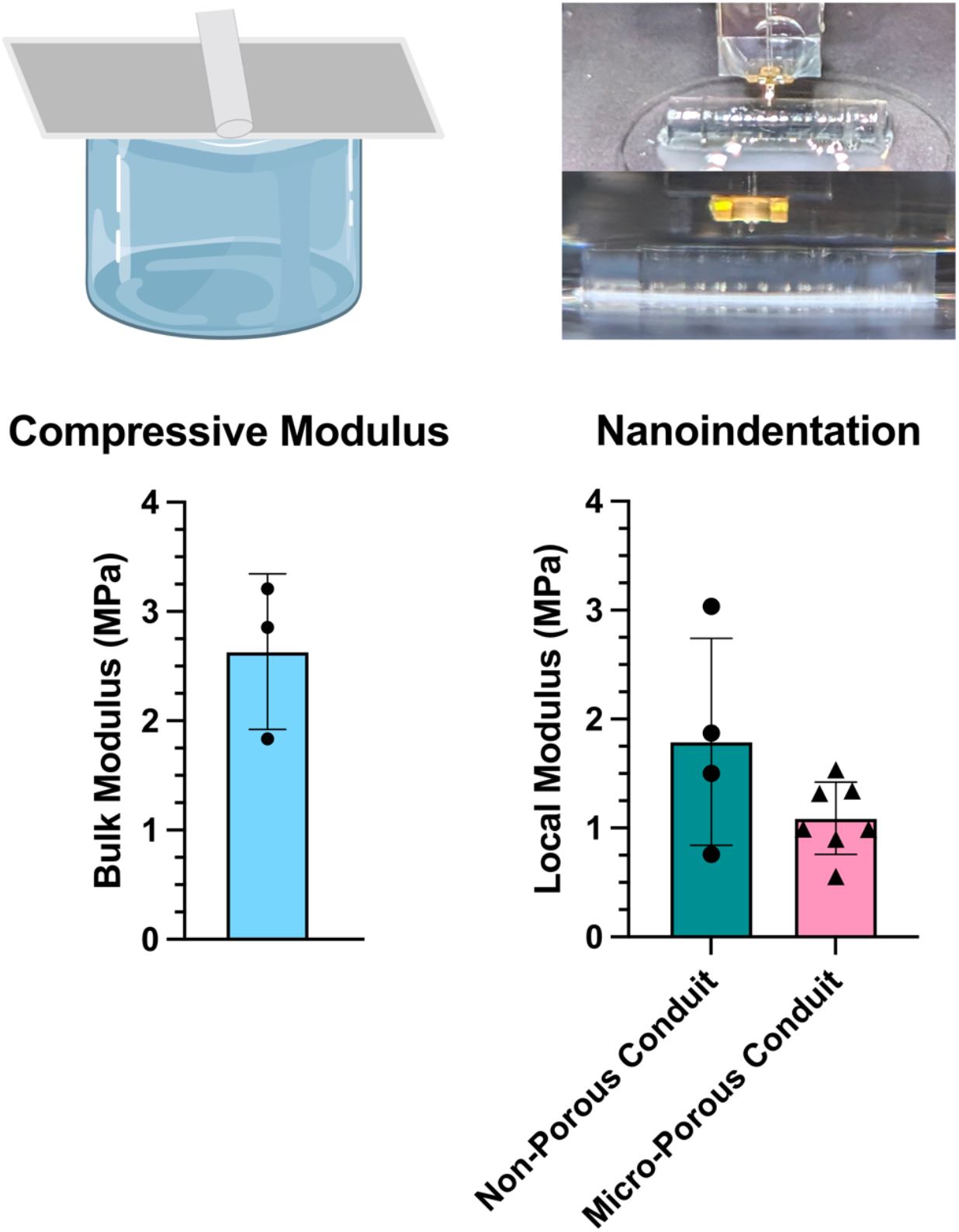
Mechanical testing. We performed both unconstrained compressive (left) and nanoindentation (right) testing to determine the bulk and local moduli, respectively. The nanoindentation was performed by sampling across the 4-mm length of the central microchannel region of the conduit. All testing were performed in a room temperature 1X DPBS environment. The nanoindentation results found no statistical difference between the local moduli of the two conduit groups. Compressive testing: n = 3. Nanoindentation testing: n = 4, n = 7. Each nanoindentation data point is an average of at least 10 independent measurements per specimen.

### Evaluation of non-porous conduit

A 12-week in vivo pilot study of the hexagonal microchannel conduit design is used to compare its performance against the clinical standard treatment, a reverse autograft. The sciatic nerve is severed at a region prior to branching and the conduit is immediately implanted and secured in place with a clinical-grade fibrin adhesive (Figure 3). Thy1-CFP transgenic mice enables the use of fluorescence in vivo microscopy to evaluate the axonal extension through the conduit microchannels in situ (Figure 3 B). Multiple fasciculated axon tracts are fully extended from the proximal end to the distal end at 12 wpi. Notably, the axons extend through the central seven microchannels as individual fascicles (Figure 1 D). We observe that the autograft results in dense external vascularization extending from the distal nerve tissue to the regenerating proximal axons (Figure 4 C). However, little to no external blood vessels extend from the distal region along the conduit (Figure 4 A). We hypothesize that the outer wall of the conduit shields the regenerating axons from the pro-regenerative external distal vasculature. We also hypothesize that introducing micropores along the conduit orthogonal to the microchannels enables crosstalk between the injury site and the distal vasculature, thus facilitating the integration of the external distal vasculature with the conduit, and subsequently the regenerating axons.

**Figure 3.**
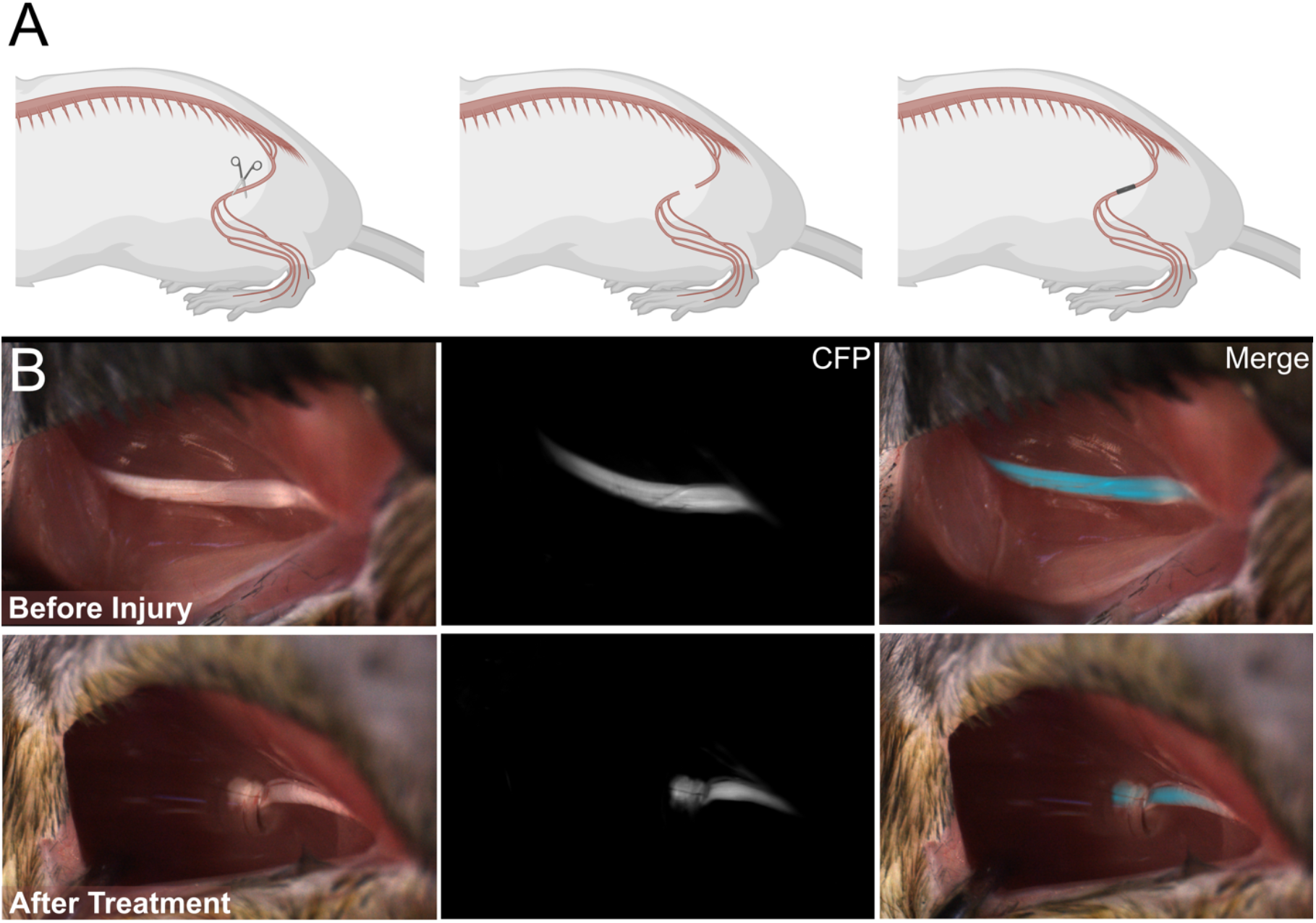
Complete transection sciatic nerve injury model and therapeutic implantation. (A) Schematic demonstrating the complete transection of the left hindlimb sciatic nerve via surgical scissors (left), retraction of the resulting two nerve stumps (middle), and implantation of conduit. The nerve stumps are secured within the conduit via a clinical-grade fibrin adhesive. (B) White light, fluorescence, and merged images taken before and after complete transection of the sciatic nerve and implantation of the nerve guidance conduit.

**Figure 4.**
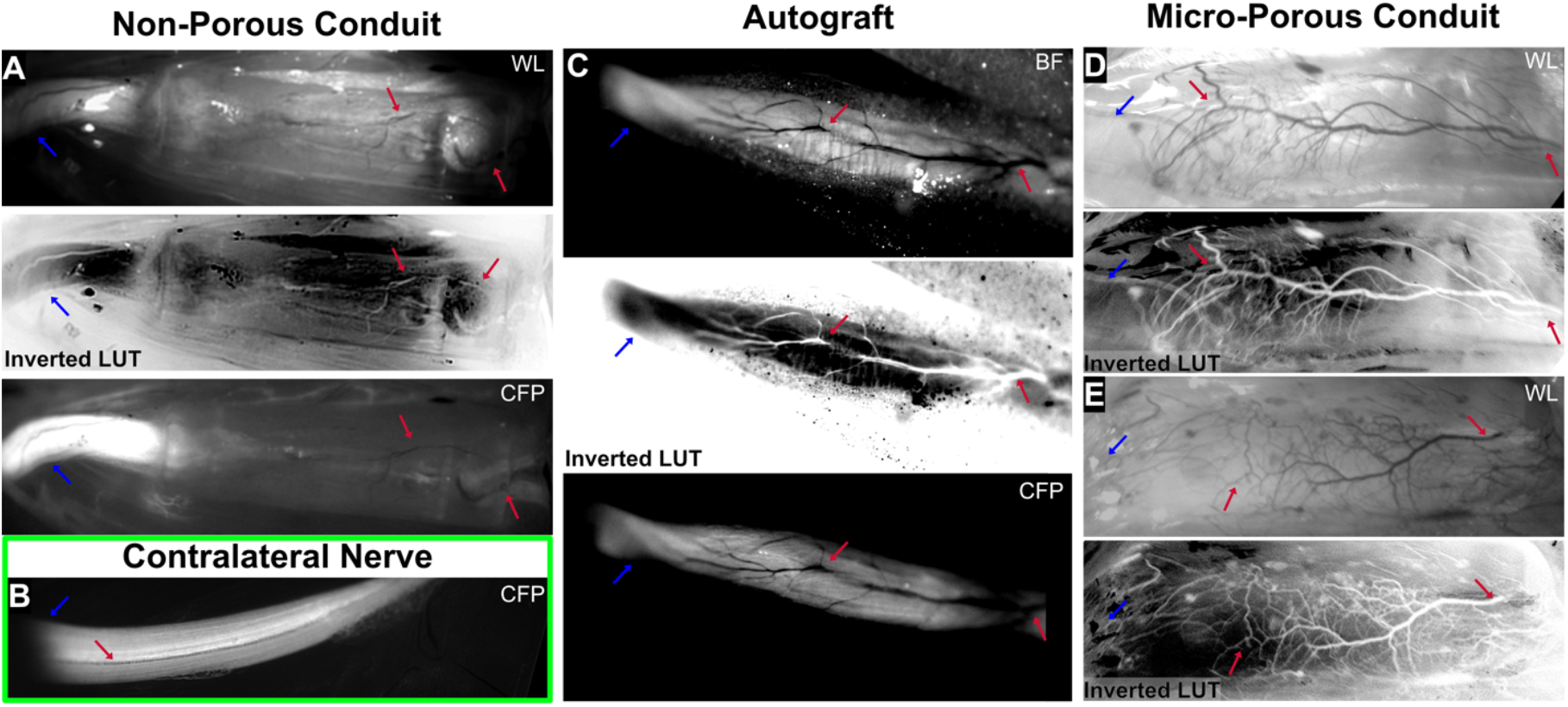
In vivo study of the therapeutic nerve guidance conduit. Two therapeutic conduit variants and a clinically employed therapeutic surgical intervention were assessed: non-porous microchannel conduit (A), reverse autologous graft (C), and non-porous microchannel conduit (D, E). White light, its corresponding inverted-color processed image, and fluorescence in vivo images taken at 12 wpi (A, C). White light in vivo images and corresponding inverted-color processed images taken at 4 wpi from two subjects (D, E). The inverted color images aid in differentiating the blood vessels from surrounding tissue and implant. Fluorescence in vivo image of a representative contralateral nerve at 17 wpi (B). Blue arrows: proximal end. Red arrows: distal extraneural vasculature growing out towards proximal regenerating axons.

### Evaluation of micro-porous conduit

A 4-week in vivo pilot study is conducted to observe if the addition of the micropores in the size and pattern as stated leads to a difference in extraneural vascularization. Indeed, distal vasculature extend along the length of the conduit and seemingly integrate with the regenerating axons (Figure 4 D, E). The morphology of the extraneural vasculature interacting with the microporous conduit is highly similar to that of the autograft, indicating that the introduction of the micropores may lead to a more physiologically guided regeneration of the severed nerve (Figure 4 C, D, E). Notably, the axon extension within the microchannels is similar to that of the non-porous conduit, which alleviates any concern that the micropores could negatively affect the aligned guidance of the regenerating axons to the distal nerve end.

### 4-month study of functional recovery

The non-porous and micro-porous conduits with the positive and negative control groups are studied over a 17-week recovery period to ensure sufficient healing time to assess functional recovery in a mouse sciatic nerve model ^41^. Dynamic gait analysis is performed longitudinally to assess the recovery of bulk motor function over time. Compound muscle action potential (CMAP) electrophysiology is performed at the end of the study. CMAP electrophysiology provides a direct assessment of the ability of the motor nerve tract to transmit an electrical signal to the muscle tissue.

To perform the electrophysiology assessment, a dual prong electrode is inserted into the gastrocnemius muscle and a stimulator probe is put in direct contact with the nerve proximal to the injury. This test is performed with the sciatic nerve exposed to ensure that the proximal nerve region was directly stimulated. We characterize the latency and peak-to-peak amplitude of the response waveform to a 0.15 mA pulse stimulation (Figure 5). Latency is the time elapsed between the stimulation pulse and the start of the action potential signal (Figure 5 A). Notably, both conduit treatment groups performed similarly to the autograft for both latency and peak-to-peak amplitude metrics; there are no significant differences between either conduit group and the autograft group (Figure 5 A, B). The two conduit and autograft treatment groups all performed better than the hollow nerve spacer negative control. Representative waveforms that most closely matched the latency and peak-to-peak amplitude means from each group are provided (Figure 5 C).

**Figure 5.**
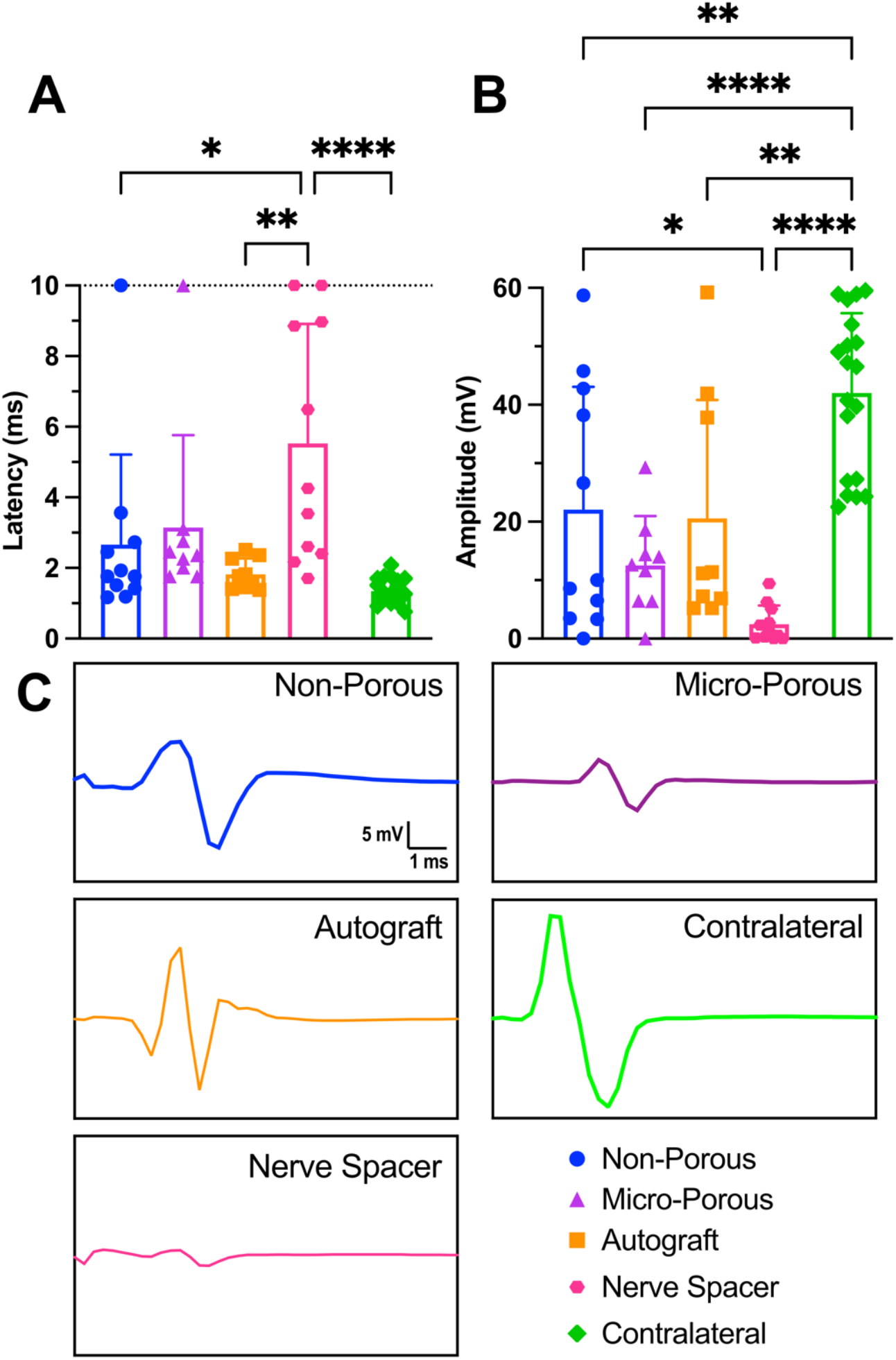
CMAP electrophysiology for assessing functional nerve regeneration. (A) Comparison of signal latency between the therapeutic conduit groups and control groups. A value of 10 ms was assigned to a waveform if no discernible signal was detected. (B) Comparison of peak-to-peak amplitude between the therapeutic conduit groups and control groups. A value of 0 mV was assigned to a waveform if no discernible signal was detected. Each data point is the average of at least 3 technical replicates; n ≥ 9. (C) Representative waveforms (i.e., closest to the mean for latency and/or amplitude) of each experimental group.

We evaluate the recovery of normal gait as an assessment of motor function by characterizing two dynamic gait parameters: left limb base of support (BOS) and left pair lag (Figure 6). Base of support is a well-established metric of stable gait ^42^. Time lags are a well-established metric of quadrupedal limb coordination during dynamic gait ^43,44^. We are interested in the left side as the injury is of the left hindlimb. The non-porous and micro-porous conduit groups both experience statistically significant changes in their gait performance in both metrics at 2 wpi. At 16 wpi, the non-porous and micro-porous conduit groups performance are both no longer statistically different than their normal gait performance prior to the injury, according to both the left limb BOS and left pair lag metrics.

**Figure 6.**
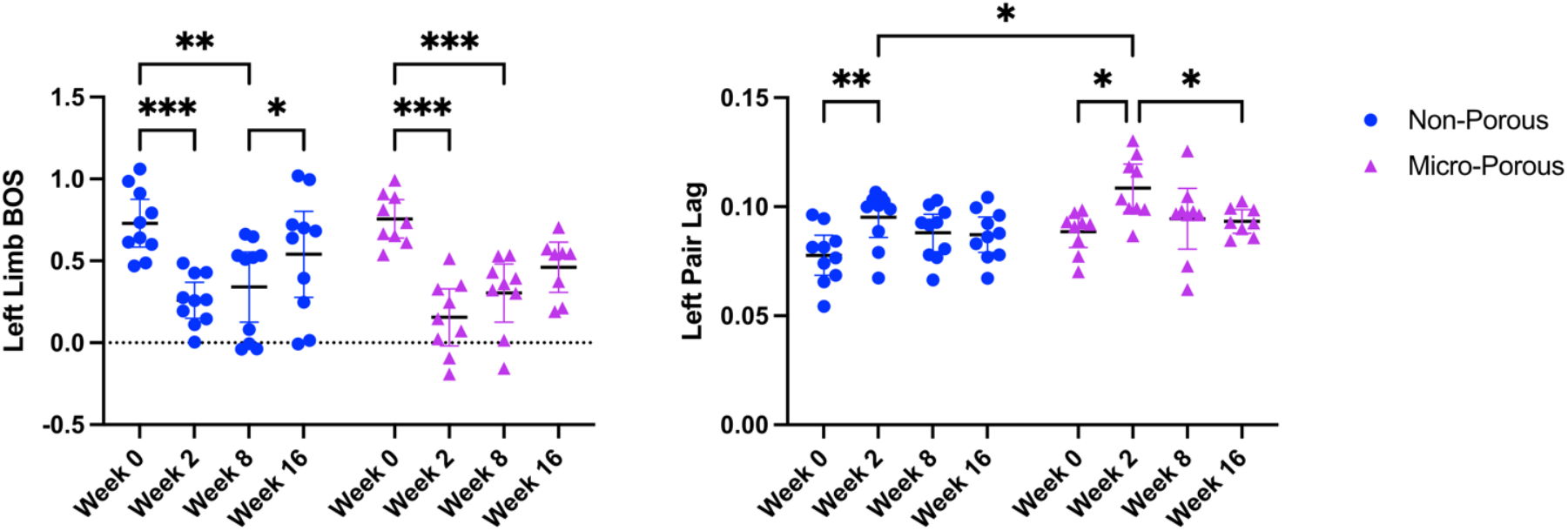
Dynamic gait analysis for assessing functional recovery after left-hindlimb sciatic nerve injury. (Left) Comparison over time of the left limb base of support (BOS) metric for the two conduit groups. (Right) Comparison over time of the left pair lag metric for the two conduit groups. * : p < 0.05; ** : p < 0.005; *** : p < 0.0005. Week 0 testing was done immediately prior to the sciatic nerve injury and conduit implantation surgery.

## Discussion

There has been much interest in incorporating bioactive components such as growth factors and cells within conduits to bolster functional regeneration ^6,45–47^. However, such an approach, even if efficacious, likely will be prohibitively expensive and will need to undergo a complex and rigorous FDA approval process as it would be considered a combination product. Ultimately, a combination device is unlikely to readily translate to the clinic to replace the need for autografts due to the cost and regulatory hurdles. Our team set out to develop a conduit whose regenerative guidance properties were solely derived from polymeric materials properties and topological cues, as such a device would have a streamlined pathway to FDA approval, and thus a high likelihood of successful clinical translation to provide therapeutic benefit to patients.

This study demonstrated that a 3D-printed PEGDA-GelMA composite hydrogel conduit with 300-μm-wide hexagonal microchannels can guide the regeneration of multiple discrete axon fascicles across a complete transection sciatic nerve injury site in a mouse model (Figure 1) and facilitate recovery of nerve function as shown by CMAP electrophysiology (Figure 5) and recovery of motor function as shown by the dynamic gait analysis (Figure 6). Demonstrating that an NGC can isolate multiple fascicles and thus could in principle guide them to different regions is a significant finding as this fulfills a critical unmet need in clinically treating complex injuries to highly branched nerves such as the facial nerves.

The RaPID 3D printing system enabled a systematic evaluation of the NGC engineering design. Upon the initial in vivo study with the non-porous conduit and autograft therapeutic conditions, we discovered that though the autograft experienced robust external vascularization from distal blood vessels (Figure 4 C), the non-porous conduit experienced minimal external distal blood vessel interaction (Figure 4 A). Angiogenesis from distal blood vessels towards injured axons is a known pro-regenerative response of the peripheral nervous system ^34^. We hypothesized that the observed difference between therapies in enabling distal angiogenesis from the extraneural blood vessels was caused by the open environment of the autograft vs the closed environment of the non-porous conduit. To test this hypothesis, we modified the conduit design by adding 50-μm-wide micropores patterned along the central microchannel region of the conduit to produce the micro-porous conduit (Figure 1 B, C). The RaPID 3D printing system readily enabled this design change. This engineering workflow is highly conducive to point-of-care manufacturing for patient-specific implants.

We observed that the micro-porous conduit led to robust angiogenesis from distal external blood vessels towards the proximal regenerating axons, similar in density and morphology to the angiogenesis observed in the autograft group (Figure 4 C, D, E). Interestingly, both the non-porous and micro-porous conduit groups resulted in similar functional nerve regeneration and full recovery of bulk motor function, via CMAP electrophysiology (Figure 5) and dynamic gait analysis (Figure 6), respectively. Notably, both NGC groups resulted in functional nerve regeneration as demonstrated by the CMAP latency and peak-to-peak amplitude metrics (Figure 5) that was not statistically different than that of the autograft group, which is the current clinical standard of surgical nerve reconstruction. The similarity between the non-porous and micro-porous NGC groups may be due to the complete transection nerve gap length of 4 mm, which is a relatively short gap distance. The pro-regenerative effect of the external distal blood vessel interaction may be more significant for longer-gap injury models in larger animals, and especially for clinical translation to humans. Thus, to better understand if there is a therapeutic improvement with the presence of micropores, a future study is needed with a larger nerve gap defect created by transecting out a multi-millimeter segment of the nerve. This is likely to be prohibitive in a mouse model due to its size ^48^.

We explored the effect of regularly patterned 50-mm-wide micropores on vascularization and functional recovery of the regenerating axons. Though the addition of the micropores resulted in the desired angiogenesis along the conduit and in the integration of blood vessels with the regenerating nerve fascicles, it did not result in an improved outcome in the functional recovery assessments when compared to the non-porous conduit. This absence of enhanced functional recovery in the presence of external blood vessel integration has also been observed by another group in rat sciatic nerve injury model with a 15-mm defect, though their conduit was a hollow tube lacking micro-topological guidance ^30^. Further studies of the microchannel conduit with varying micropore sizes, densities, and spatial patterns will be required to understand the role of angiogenesis from the distal external vasculature in sciatic nerve regeneration with respect to functional recovery in larger animal models.

## MATERIALS AND METHODS

### Study design

This study aimed to investigate how microscale topology can effectively guide functional neural regeneration without the aid of biologics. We previously have developed the RaPID 3D-printing system to produce nerve guidance conduits with four 400-μm-diameter channels ^39^. In that study, we demonstrated the biocompatibility and axonal-guidance efficacy of the PEGDA-GelMA hydrogel nerve conduit in a mouse sciatic nerve defect model. Based on this previous work, we hypothesized that reducing the microchannel diameter would maximize the contact area between the regenerating axon bundle and microchannel. We also hypothesized that adding regularly patterned micropores along the length of the conduit would improve microvascular integration. The designed conduit of this study had nineteen (19) microchannels with hexagonal cross-sections for increased channel density and inscribed in a 300-μm-diameter circle. To maximize the number of microchannels and reduce the spacing between regenerating nerve fascicles, the microchannel wall thickness was 10 μm. The nerve guidance conduit was specifically designed for a young-adult mouse sciatic nerve complete transection injury model which results in a 4-mm gap defect. The total length of the conduit was approximately 6 mm, accounting for the approximately 1-mm-long hallow sleeves on each end for securing the severed nerve endings via clinical-grade fibrin glue. Two therapeutic conduits, the non-porous and micro-porous microchannel conduits, were compared against a reverse-polarity 2-mm-long autograft and a 3D-printed hollow nerve spacer (negative control). We used only male mice for this study to minimize the statistical variance as this was a first in kind study comparing these novel conduit designs.

### RaPID printing of conduits

Gelatin methacryloyl (GelMA), 85% degree of methacryloylation, was synthesized as previously reported from gelatin Type A, bloom 300 (Sigma Aldrich) ^51,52^. Poly (ethylene glycol) acrylate (PEGDA, M_n_ = 700 Da) was purchased from Millipore-Sigma. Lithium phenyl(2,4,6-trimethylbenzoyl)phosphinate (LAP) was purchased from TCI America. Tartrazine was purchased from Sigma-Aldrich. The development of the rapid projection, image-guided, direct-printing (RaPID) 3D printing system was detailed in our prior work ^39^. All concentrations stated as percentages are (*w/v*)% unless explicitly stated otherwise. The prepolymer solution was composed of 25% PEGDA, 7.5% GelMA, 1% LAP, and 0.075% tartrazine in 1x Dulbecco’s phosphate-buffered saline (DPBS) solution. PEGDA provides physiological stiffness matching to native nerve tissue, GelMA provides integrin binding domains to facilitate cell attachment as it is derived from collagen, LAP serves as a cytocompatible water-soluble photoinitiator, and tartrazine is a nontoxic water-soluble photoabsorber ^53^. A photoabsorber, tartrazine, was included in the prepolymer solution to produce conduits with the 10-μm channel wall thickness.

The conduits were designed using CAD software (AutoCAD, Autodesk, Inc.), exported as an STL file, and sliced into a set of digital 2D images via a custom MATLAB script. The dimensions for the conduit design variants are 6 mm in total length, 1-mm in sleeve length, 1.75-mm outer diameter, and 1.25-mm inner diameter. The treatment conduit variants have hexagonal microchannels defined by a long diagonal length of 300 μm and channel wall thickness of 10 μm. The microchannels are 4-mm long to match the gap distance that results from a complete transection injury of the mouse sciatic nerve. The micro-porous conduit has a regularly patterned array of square micropores with a 50-μm side length spaced apart 250 μm longitudinally and circumferentially, orthogonal to the longitudinal microchannels. A negative control version of the conduit design was made by eliminating all but the two sleeve ends separated the same 4-mm distance apart with four equally spaced columns, each having cross-sectional dimensions of approximately 0.4 mm by 0.25 mm.

### Mechanical testing

The bulk Young’s modulus of the formulated hydrogel material was characterized via an unconfined uniaxial compressive test using the Univert mechanical tester (CellScale, Canada). Cylindrical specimens were prepared with a diameter of 500 μm and a height of 500 μm. The Young’s modulus was calculated from a linear regression analysis between 5% and 15% strain. *n* = 3.

The local Young’s modulus of each microchannel conduit design was characterized via nanoindentation (Piuma, Optics11 Life, Netherlands). For each specimen, a full-length conduit was fabricated to the same specifications as the in vivo implants. The conduit was secured to a plastic Petri dish using a small dot of a commercial cyanoacrylate adhesive (Loctite 401), and the conduit was then fully immersed in 1X DPBS solution. The nanoindenter probe tip was 58 μm in diameter, and the cantilever had a spring constant *k* of 4.580 N/m. A minimum of 10 (and a maximum of 30) independent measurements were made along each conduit. Non-porous conduit: *n* = 4. Micro-porous conduit: *n* = 7. The force-displacement curves were analyzed using the Hertzian model for hemispherical samples in the DataViewer software V2.6.0 (Optics 11 Life). The Poisson’s ratio was assumed to be 0.5.

### Surgical procedures

We used male BL/6 mice purchased from Charles River Labs for studying the two therapeutic conduit variants. We used male transgenic Thy1-CFP BL/6 mice (The Jackson Laboratories, US) from inbred colonies for all other *in vivo* studies (including pilot studies of the two therapeutic conduit variants). All mice were between 15-18 weeks in age at the date of surgery. The transgenic mice expressed cyan fluorescence protein (CFP) in nerve tissue, allowing for *in situ* visualization of nerve regeneration. The mice were housed in a pathogen-free vivarium maintained by the University of California San Diego Animal Care Program. All animal procedures were reviewed and approved by the University of California San Diego Institutional Animal Care and Use Committee (IACUC) to ensure compliance with all tenets of the Animal Welfare Act and PHS Policy (Protocol #S04011). All surgeries were conducted under isoflurane anesthesia (3% induction, 1-3% maintenance). After surgical intervention, the incision site was sutured closed, and the analgesic buprenorphine was subcutaneously administered. No more than four mice were kept in the same cage.

### Motor function analysis

Dynamic gait analysis was performed on the GaitLab (ViewPoint Life Sciences, France). A single researcher performed all related mice handling, and they were blinded to the experimental groups during behavioral testing and data analysis. The mice were trained on the walking platform one to two times a day for one to two weeks before surgery. Training involved allowing the mice to familiarize themselves with the GaitLab platform and running across the walkway with a consistent gait at maximum speed multiple times per session. Healthy cohort data for analysis was collected at the end of training, prior to surgery. The two treatment conduit groups (n = 10) were tested once every 2 weeks, starting two weeks after surgery for sixteen (16) weeks.

### Electrophysiology

Compound muscle action potential (CMAP) electrophysiology was performed seventeen weeks postinjury, before the nerve specimens were harvested, in both the ipsilateral and contralateral sciatic nerves. Electrophysiology was performed using a NIM-Response 2.0 nerve integrity monitor system (Medtronic, US). The recording electrode was inserted into the gastrocnemius muscle, and a ground electrode was inserted subcutaneously. The stimulating electrode was applied to the proximal nerve and delivered an electrical stimulus of 10-15 mA at 1 pulse/s with 0.1 ms duration. A minimum of three replicate waveforms were obtained per nerve and averaged. The averaged waveforms were analyzed for latency and peak-to-peak amplitude based upon prior reported protocols.^54,55^

### Statistical Analysis

Statistical analysis was performed using GraphPad Prism (GraphPad Software, Inc.) and G*Power v3.1.9.6.^56^ An a priori power analysis test was performed based on preliminary mouse behavioral testing data (effect size *f* was 0.948) to determine the minimum group size needed to achieve a power of 0.8 and an *α* error probability of 0.05. The result of which was a group size of *n* = 5, which resulted in an actual power of 0.896. Descriptive statistics and their representation in figures were presented as mean ± standard deviation. Comparisons between two independent groups were conducted using Welch’s unpaired *t*-test. Outliers, when identified amongst repeated measurements, were removed using the ROUT method where Q = 1%. For comparing multiple groups, an ordinary one-way ANOVA was performed. For comparing multiple groups over time, either a repeated measures two-way ANOVA or mixed-effects model with the Geisser-Greenhouse correction was performed. The two-way ANOVA analysis cannot handle missing values, so a mixed-effects model was used in its stead when missing values were present. All ad hoc multiple comparisons testing was performed using Šídák’s with a 95% confidence interval.

## CONCLUSIONS

Overall, this study has demonstrated that in a mouse complete transection sciatic nerve injury model a RaPID 3D-printed PEGDA-GelMA composite hydrogel with 300-μm-wide hexagonal microchannels with 10-μm channel thickness leads to functional axonal regeneration after a 4-month recovery time period that is quantitatively similar in functional nerve regeneration performance to that of the clinically used reverse autograft surgical therapy and results in restoration of bulk motor function to pre-injury performance. This finding is remarkable in that the therapeutic conduit contained no xenogeneic, bioactive, or cellular components. Additionally, the technology behind the RaPID bioprinting system has been successfully commercially translated^49,50^. We are currently working towards obtaining investigational device exemption (IDE) approval from the FDA. In parallel, we also plan to apply for the FDA Early Feasibility Study (EFS) program, which would enable us to perform a small pilot study in human patients.

## Acknowledgements

The authors would like to thank Henry H. Hwang for assistance with the Autodesk AutoCAD software, and Michael Whitney for assistance in study design considerations and in initial discussions that led to the conceptualization of the hollow nerve spacer. Thank you to Shangting You for developing the STL slicer MATLAB script. The authors would like to thank the University of California San Diego School of Medicine Microscopy Core, which is supported by the NIH grant NINDS P30NS047101, for access to their confocal microscopes. The authors would like to thank Laarni Gapuz and Kimberly Mcintyre, of the University of California San Diego Biorepository and Tissue Technology Shared Resources, for assistance in optimizing the nerve conduit sectioning protocol. This material is based upon work supported by the National Science Foundation Graduate Research Fellowship under Grant No. DGE-1650112 (J.S.). J.S. acknowledges funding support by the National Institute of Neurological Disorders and Stroke (NINDS) of the National Institutes of Health under Award Number F31NS125986. Research reported in this publication was supported by the Eunice Kennedy Shriver National Institute of Child Health and Human Development (NICHD) of the National Institutes of Health under award numbers HD090662 and HD112026. The content is solely the responsibility of the authors and does not necessarily represent the official views of the National Institutes of Health.

## Data Sharing

The data that support the findings of this study are available from the corresponding authors upon reasonable request.

## References

1. Kasper M, Deister C, Beck F, Schmidt CE. Bench-to-Bedside Lessons Learned: Commercialization of an Acellular Nerve Graft. Adv Healthcare Materials. 2020;9(16):2000174. doi:10.1002/adhm.202000174

2. Millesi H. Bridging defects: autologous nerve grafts. In: Millesi H, Schmidhammer R, eds. How to Improve the Results of Peripheral Nerve Surgery. Acta Neurochirurgica Supplementum. Springer; 2007:37–38. doi:10.1007/978-3-211-72958-8_8

3. Grinsell D, Keating CP. Peripheral Nerve Reconstruction after Injury: A Review of Clinical and Experimental Therapies. BioMed Research International. 2014;2014:e698256. doi:10.1155/2014/698256

4. Ducic I, Yoon J, Buncke G. Chronic postoperative complications and donor site morbidity after sural nerve autograft harvest or biopsy. Microsurgery. 2020;40(6):710–716. doi:10.1002/micr.30588

5. Hudson TW, Evans GRD, Schmidt CE. ENGINEERING STRATEGIES FOR PERIPHERAL NERVE REPAIR. Orthopedic Clinics of North America. 2000;31(3):485–497. doi:10.1016/S0030-5898(05)70166-8

6. Lackington WA, Ryan AJ, O’Brien FJ. Advances in Nerve Guidance Conduit-Based Therapeutics for Peripheral Nerve Repair. ACS Biomater Sci Eng. 2017;3(7):1221–1235. doi:10.1021/acsbiomaterials.6b00500

7. Sun AX, Prest TA, Fowler JR, Brick RM, Gloss KM, Li X, DeHart M, Shen H, Yang G, Brown BN, Alexander PG, Tuan RS. Conduits harnessing spatially controlled cell-secreted neurotrophic factors improve peripheral nerve regeneration. Biomaterials. 2019;203:86–95. doi:10.1016/j.biomaterials.2019.01.038

8. Chen BK, Madigan NN, Hakim JS, Dadsetan M, McMahon SS, Yaszemski MJ, Windebank AJ. GDNF Schwann cells in hydrogel scaffolds promote regional axon regeneration, remyelination and functional improvement after spinal cord transection in rats. Journal of Tissue Engineering and Regenerative Medicine. 2018;12(1):e398–e407. doi:10.1002/term.2431

9. Adidharma W, Wang Y, Kotsis SV, Wang L, Chung KC. Utilization Trends of Nerve Autograft Alternatives for the… : Plastic and Reconstructive Surgery. Accessed February 4, 2025. https://journals.lww.com/plasreconsurg/fulltext/2024/04000/utilization_trends_of_nerve_autograft_alternatives.24.aspx

10. Pabari A, Lloyd-Hughes H, Seifalian AM, Mosahebi A. Nerve Conduits for Peripheral Nerve Surgery. Plastic and Reconstructive Surgery. 2014;133(6):1420. doi:10.1097/PRS.0000000000000226

11. Song J, Dong J, Yuan Z, Huang M, Yu X, Zhao Y, Shen Y, Wu J, EL-Newehy M, Abdulhameed MM, Sun B, Chen J, Mo X. Shape-Persistent Conductive Nerve Guidance Conduits for Peripheral Nerve Regeneration. Advanced Healthcare Materials. 2024;13(26):2401160. doi:10.1002/adhm.202401160

12. Ramesh PA, Sethuraman S, Subramanian A. Fabrication of Anatomically Equivalent Pectin-Based Multifilament Nerve Conduits. ACS Appl Bio Mater. 2024;7(10):6706–6719. doi:10.1021/acsabm.4c00872

13. Belanger K, Schlatter G, Hébraud A, Marin F, Testelin S, Dakpé S, Devauchelle B, Egles C. A multi-layered nerve guidance conduit design adapted to facilitate surgical implantation. Health Science Reports. 2018;1(12):e86. doi:10.1002/hsr2.86

14. Carvalho CR, Oliveira JM, Reis RL. Modern Trends for Peripheral Nerve Repair and Regeneration: Beyond the Hollow Nerve Guidance Conduit. Front Bioeng Biotechnol. 2019;7. doi:10.3389/fbioe.2019.00337

15. Wang J, Xiong H, Zhu T, Liu Y, Pan H, Fan C, Zhao X, Lu WW. Bioinspired Multichannel Nerve Guidance Conduit Based on Shape Memory Nanofibers for Potential Application in Peripheral Nerve Repair. ACS Nano. 2020;14(10):12579–12595. doi:10.1021/acsnano.0c03570

16. Hwangbo H, Kim W, Kim GH. Lotus-Root-Like Microchanneled Collagen Scaffold. ACS Appl Mater Interfaces. Published online December 2, 2020. doi:10.1021/acsami.0c14670

17. Wang J, Cheng Y, Wang H, Wang Y, Zhang K, Fan C, Wang H, Mo X. Biomimetic and Hierarchical Nerve Conduits from Multifunctional Nanofibers for Guided Peripheral Nerve Regeneration. Acta Biomaterialia. Published online September 2020:S1742706120305602. doi:10.1016/j.actbio.2020.09.037

18. Bozkurt A, Lassner F, O’Dey D, Deumens R, Böcker A, Schwendt T, Janzen C, Suschek CV, Tolba R, Kobayashi E, Sellhaus B, Tholl S, Eummelen L, Schügner F, Olde Damink L, Weis J, Brook GA, Pallua N. The role of microstructured and interconnected pore channels in a collagen-based nerve guide on axonal regeneration in peripheral nerves. Biomaterials. 2012;33(5):1363–1375. doi:10.1016/j.biomaterials.2011.10.069

19. Belkas JS, Shoichet MS, Midha R. Axonal guidance channels in peripheral nerve regeneration. Operative Techniques in Orthopaedics. 2004;14(3):190–198. doi:10.1053/j.oto.2004.06.001

20. Stokols S, Sakamoto J, Breckon C, Holt T, Weiss J, Tuszynski MH. Templated Agarose Scaffolds Support Linear Axonal Regeneration. Tissue Engineering. 2006;12(10):2777–2787. doi:10.1089/ten.2006.12.2777

21. Bellamkonda RV. Peripheral nerve regeneration: An opinion on channels, scaffolds and anisotropy. Biomaterials. 2006;27(19):3515–3518. doi:10.1016/j.biomaterials.2006.02.030

22. Sun X, Bai Y, Zhai H, Liu S, Zhang C, Xu Y, Zou J, Wang T, Chen S, Zhu Q, Liu X, Mao H, Quan D. Devising micro/nano-architectures in multi-channel nerve conduits towards a pro-regenerative matrix for the repair of spinal cord injury. Acta Biomaterialia. 2019;86:194–206. doi:10.1016/j.actbio.2018.12.032

23. Zeng C guang, Xiong Y, Xie G, Dong P, Quan D. Fabrication and Evaluation of PLLA Multichannel Conduits with Nanofibrous Microstructure for the Differentiation of NSCs In Vitro. Tissue Engineering Part A. 2013;20(5-6):1038–1048. doi:10.1089/ten.tea.2013.0277

24. Pawelec KM, Koffler J, Shahriari D, Galvan A, Tuszynski MH, Sakamoto J. Microstructure and in vivo characterization of multi-channel nerve guidance scaffolds. Biomed Mater. 2018;13(4):044104. doi:10.1088/1748-605X/aaad85

25. Shahriari D, Shibayama M, Lynam DA, Wolf KJ, Kubota G, Koffler JY, Tuszynski MH, Campana WM, Sakamoto JS. Peripheral nerve growth within a hydrogel microchannel scaffold supported by a kink-resistant conduit. Journal of Biomedical Materials Research Part A. 2017;105(12):3392–3399. doi:10.1002/jbm.a.36186

26. Dinis TM, Elia R, Vidal G, Dermigny Q, Denoeud C, Kaplan DL, Egles C, Marin F. 3D multi-channel bi-functionalized silk electrospun conduits for peripheral nerve regeneration. Journal of the Mechanical Behavior of Biomedical Materials. 2015;41:43–55. doi:10.1016/j.jmbbm.2014.09.029

27. Tu X, Wang L, Wei J, Wang B, Tang Y, Shi J, Zhang Z, Chen Y. 3D printed PEGDA microstructures for gelatin scaffold integration and neuron differentiation. Microelectronic Engineering. 2016;158:30–34. doi:10.1016/j.mee.2016.03.007

28. Zhu W, Tringale KR, Woller SA, You S, Johnson S, Shen H, Schimelman J, Whitney M, Steinauer J, Xu W, Yaksh TL, Nguyen QT, Chen S. Rapid continuous 3D printing of customizable peripheral nerve guidance conduits. Materials Today. 2018;21(9):951–959. doi:10.1016/j.mattod.2018.04.001

29. Moimas S, Novati F, Ronchi G, Zacchigna S, Fregnan F, Zentilin L, Papa G, Giacca M, Geuna S, Perroteau I, Arnež ZM, Raimondo S. Effect of vascular endothelial growth factor gene therapy on post-traumatic peripheral nerve regeneration and denervation-related muscle atrophy. Gene Ther. 2013;20(10):1014–1021. doi:10.1038/gt.2013.26

30. Hromada C, Heimel P, Kerbl M, Gál L, Nürnberger S, Schaedl B, Ferguson J, Swiadek N, Monforte X, Heinzel JC, Nógrádi A, Teuschl-Woller AH, Hercher D. Silk-based nerve guidance conduits with macroscopic holes modulate the vascularization of regenerating rat sciatic nerve. Neural Regeneration Research. Published online May 13, 2024. doi:10.4103/NRR.NRR-D-23-01518

31. Huang Y, Ye K, He A, Wan S, Wu M, Hu D, Xu K, Wei P, Yin J. Dual-layer conduit containing VEGF-A – Transfected Schwann cells promotes peripheral nerve regeneration via angiogenesis. Acta Biomaterialia. Published online March 30, 2024. doi:10.1016/j.actbio.2024.03.029

32. Caillaud M, Richard L, Vallat JM, Desmoulière A, Billet F. Peripheral nerve regeneration and intraneural revascularization. Neural Regeneration Research. 2019;14(1):24. doi:10.4103/1673-5374.243699

33. Wu P, Tong Z, Luo L, Zhao Y, Chen F, Li Y, Huselstein C, Ye Q, Ye Q, Chen Y. Comprehensive strategy of conduit guidance combined with VEGF producing Schwann cells accelerates peripheral nerve repair. Bioactive Materials. 2021;6(10):3515–3527. doi:10.1016/j.bioactmat.2021.03.020

34. Cattin AL, Burden JJ, Van Emmenis L, Mackenzie FE, Hoving JJA, Garcia Calavia N, Guo Y, McLaughlin M, Rosenberg LH, Quereda V, Jamecna D, Napoli I, Parrinello S, Enver T, Ruhrberg C, Lloyd AC. Macrophage-Induced Blood Vessels Guide Schwann Cell-Mediated Regeneration of Peripheral Nerves. Cell. 2015;162(5):1127–1139. doi:10.1016/j.cell.2015.07.021

35. Bhise MG, Patel L, Patel K. 3D Printed Medical Devices: Regulatory Standards and Technological Advancements in the USA, Canada and Singapore-A Cross-National Study. Int J Pharm Investigation. 2025;14(3):888–902. doi:10.5530/ijpi.14.3.99

36. Discussion Paper: 3D Printing Medical Devices at the Point of Care. FDA; 2021. https://www.regulations.gov/document/FDA-2021-N-1272-0001

37. Wood LB, Ge R, Kamm RD, Asada HH. Nascent vessel elongation rate is inversely related to diameter in in vitro angiogenesis. Integrative Biology. 2012;4(9):1081–1089. doi:10.1039/c2ib20054f

38. Schimelman J. Created in Biorender.; 2025. https://BioRender.com/l51n186

39. Zhu W, Tringale KR, Woller SA, You S, Johnson S, Shen H, Schimelman J, Whitney M, Steinauer J, Xu W, Yaksh TL, Nguyen QT, Chen S. Rapid continuous 3D printing of customizable peripheral nerve guidance conduits. Materials Today. 2018;21(9):951–959. doi:10.1016/j.mattod.2018.04.001

40. Petit E, Bavykina V, Thibault M, Bilodeau A, Choinière W, Brosseau JP, Laurent B, Lauzon MA. Assessing tissue mechanical properties: Development of a custom-made tensile device and application on rodents sciatic nerves. Journal of the Mechanical Behavior of Biomedical Materials. 2024;159:106709. doi:10.1016/j.jmbbm.2024.106709

41. Saller MM, Huettl RE, Mayer JM, Feuchtinger A, Krug C, Holzbach T, Volkmer E. Validation of a novel animal model for sciatic nerve repair with an adipose-derived stem cell loaded fibrin conduit. Neural Regen Res. 2018;13(5):854–861. doi:10.4103/1673-5374.232481

42. Lugade V, Lin V, Chou LS. Center of mass and base of support interaction during gait. Gait & Posture. 2011;33(3):406–411. doi:10.1016/j.gaitpost.2010.12.013

43. Abourachid A. A new way of analysing symmetrical and asymmetrical gaits in quadrupeds. Comptes Rendus Biologies. 2003;326(7):625–630. doi:10.1016/S1631-0691(03)00170-7

44. Maes L, Abourachid A. Gait transitions and modular organization of mammal locomotion. Journal of Experimental Biology. 2013;216(12):2257–2265. doi:10.1242/jeb.082149

45. Kuffler DP, Foy C. Restoration of Neurological Function Following Peripheral Nerve Trauma. International Journal of Molecular Sciences. 2020;21(5):1808. doi:10.3390/ijms21051808

46. Alrashdan MS, Sung MA, Kim Kwon Y, Chung HJ, Kim SJ, Lee JH. Effects of combining electrical stimulation with BDNF gene transfer on the regeneration of crushed rat sciatic nerve. Acta Neurochir. 2011;153(10):2021–2029. doi:10.1007/s00701-011-1054-x

47. Yeh CW, Wang LW, Wu HC, Hsieh YK, Wang J, Chen MH, Wang TW. Development of biomimetic micro-patterned device incorporated with neurotrophic gradient and supportive Schwann cells for the applications in neural tissue engineering. Biofabrication. 2017;9(1):015024. doi:10.1088/1758-5090/aa5ef2

48. Angius D, Wang H, Spinner RJ, Gutierrez-Cotto Y, Yaszemski MJ, Windebank AJ. A systematic review of animal models used to study nerve regeneration in tissue-engineered scaffolds. Biomaterials. 2012;33(32):8034–8039. doi:10.1016/j.biomaterials.2012.07.056

49. Listek V. BICO Buys Bioprinting Startup Allegro 3D—8th Acquisition in a Year. 3DPrint.com | The Voice of 3D Printing / Additive Manufacturing. May 10, 2022. Accessed March 20, 2025. https://3dprint.com/291132/bico-buys-bioprinting-startup-allegro-3d-8th-acquisition-in-a-year/

50. Bionova X. CELLINK. Accessed March 20, 2025. https://www.cellink.com/bioprinting/bionova-x/

51. Zhong Z, Deng X, Wang P, Yu C, Kiratitanaporn W, Wu X, Schimelman J, Tang M, Balayan A, Yao E, Tian J, Chen L, Zhang K, Chen S. Rapid bioprinting of conjunctival stem cell micro-constructs for subconjunctival ocular injection. Biomaterials. 2021;267:120462. doi:10.1016/j.biomaterials.2020.120462

52. Shirahama H, Lee BH, Tan LP, Cho NJ. Precise Tuning of Facile One-Pot Gelatin Methacryloyl (GelMA) Synthesis. Scientific Reports. 2016;6:srep31036. doi:10.1038/srep31036

53. Grigoryan B, Paulsen SJ, Corbett DC, Sazer DW, Fortin CL, Zaita AJ, Greenfield PT, Calafat NJ, Gounley JP, Ta AH, Johansson F, Randles A, Rosenkrantz JE, Louis-Rosenberg JD, Galie PA, Stevens KR, Miller JS. Multivascular networks and functional intravascular topologies within biocompatible hydrogels. Science. 2019;364(6439):458–464. doi:10.1126/science.aav9750

54. Pollari E, Prior R, Robberecht W, Van Damme P, Van Den Bosch L. In Vivo Electrophysiological Measurement of Compound Muscle Action Potential from the Forelimbs in Mouse Models of Motor Neuron Degeneration. J Vis Exp. 2018;(136). doi:10.3791/57741

55. Arnold WD, Sheth KA, Wier CG, Kissel JT, Burghes AH, Kolb SJ. Electrophysiological Motor Unit Number Estimation (MUNE) Measuring Compound Muscle Action Potential (CMAP) in Mouse Hindlimb Muscles. JoVE (Journal of Visualized Experiments). 2015;(103):e52899. doi:10.3791/52899

56. Faul F, Erdfelder E, Lang AG, Buchner A. G*Power 3: A flexible statistical power analysis program for the social, behavioral, and biomedical sciences. Behavior Research Methods. 2007;39(2):175–191. doi:10.3758/BF03193146

